## Supplementary material for "Lumbar corticospinal tract in rodents modulates sensory inputs but does not convey motor command": Suppl. Figures

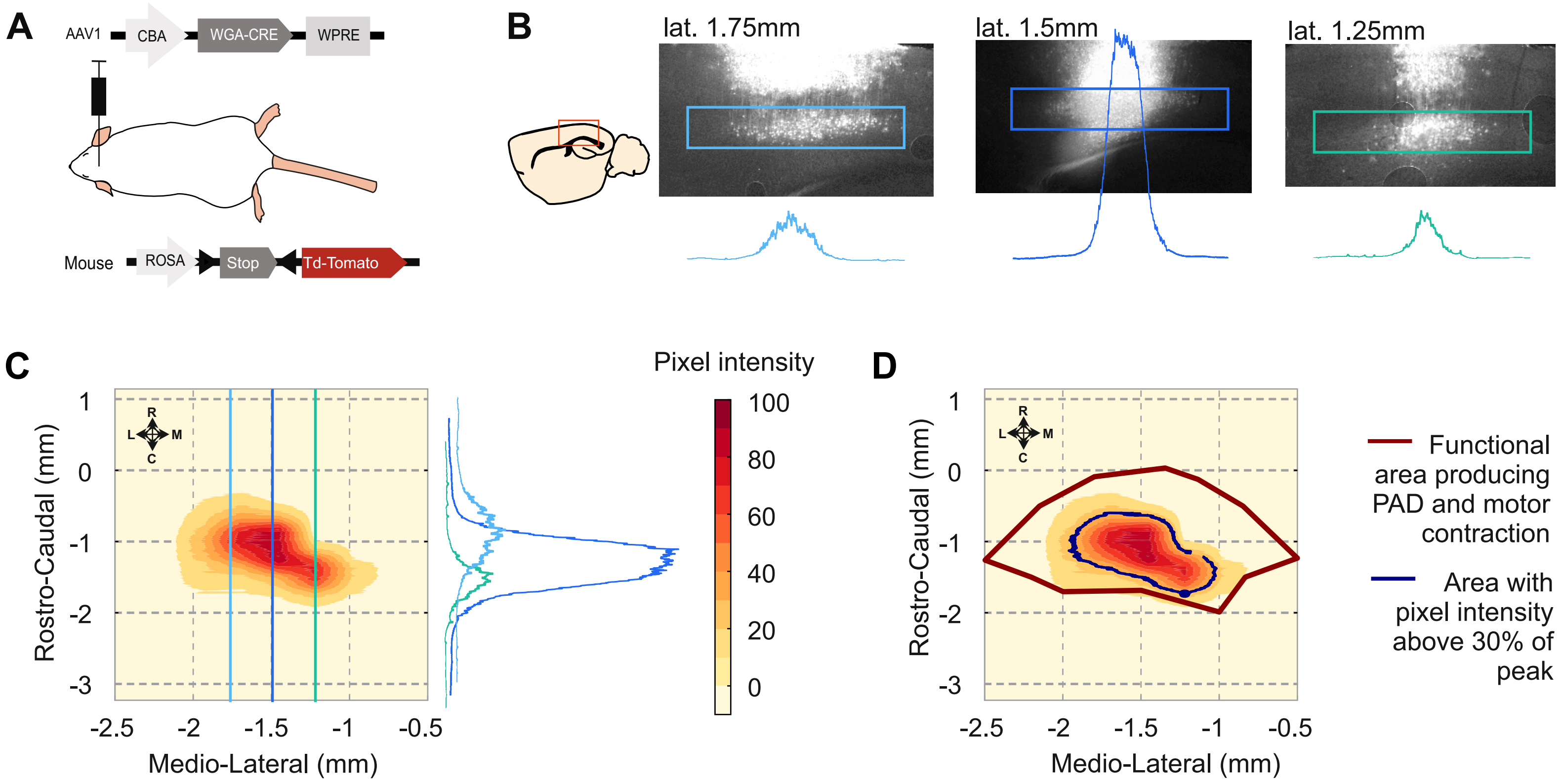


**Figure 1 - Figure Supplement 1. Sensorimotor injection site analyses : cortical layer V fluorescence quantification.**
A. Schematic experimental design to transynaptically label cortical postsynaptic neurons.
B. *Left*, drawing of a sagittal slice of the mouse brain illustrating (red square) the localization of the pictures at the right. *Right*, images of the sensorimotor cortex of a single animal, at different lateral positions from the midline. The blue squares show the analyzed area, comprising cortical layer V. The plots show the vertical averages of pixel intensity throughout each blue square. The 3 slices are localized in the sites indicated by the blue lines in C.
C. Example of an injection site represented as a heat map of the pixel intensity across the layer V of a cortex. The map is build using the three plots presented in B (illustrated here in the right to show their orientation) as well as additional ones from sequential brain sections on the same animal.
D. Overlap between the heat map in C and the functional area eliciting motor contraction and PAD (Red line). The black line indicates the area with a pixel intensity above 30 %.

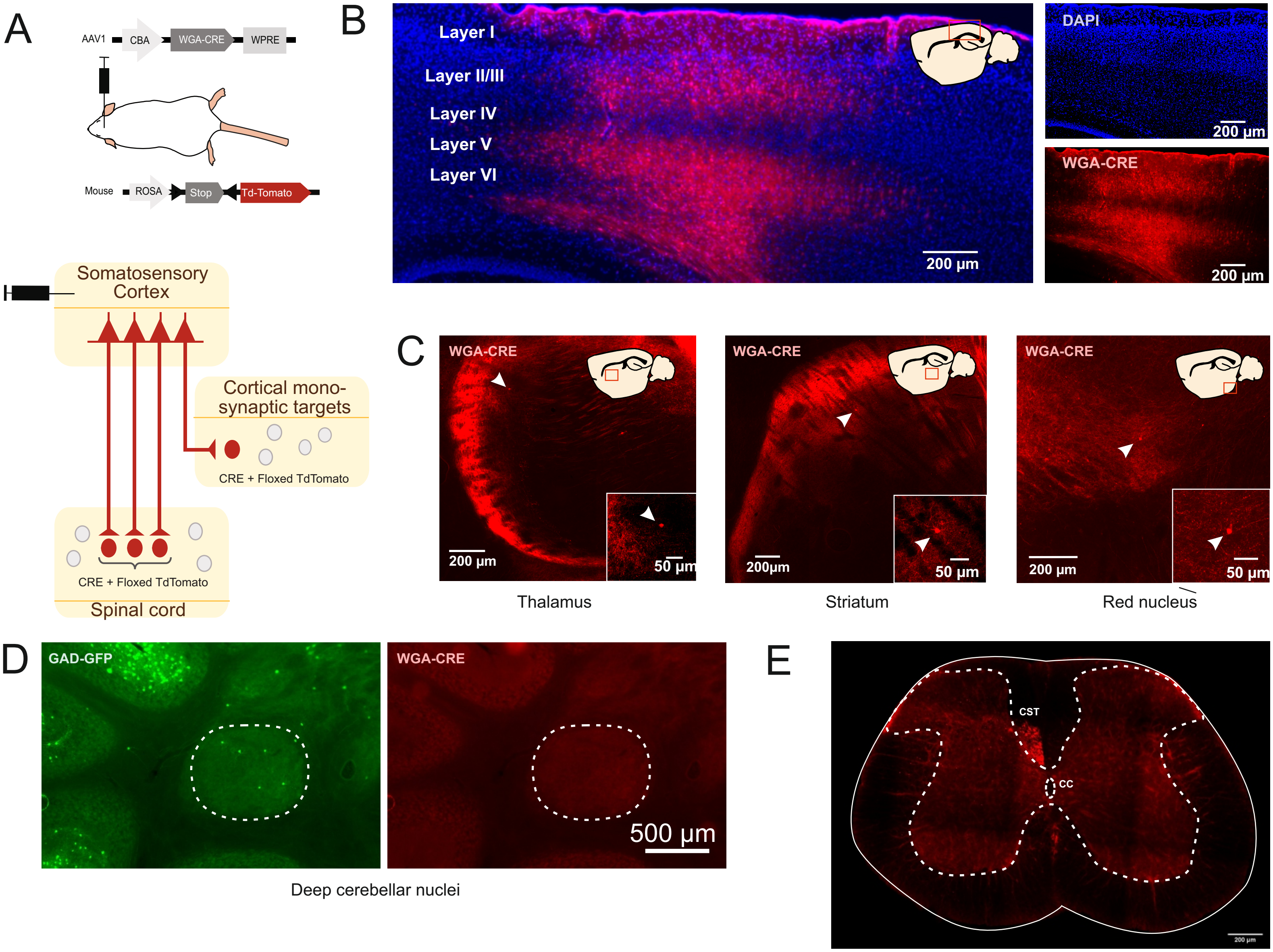


**Figure 1 - Figure Supplement 2. Monosynaptic transynaptic tracing from the cortex**

A. Schematic experimental design to transynaptically label cortical postsynaptic neurons.

B. Sagittal image of the injection site, in the sensorimotor cortex, illustrating TdTomato+ neurons (red) and DAPI (blue). The drawing inset shows the area corresponding to the image.

C. Example of stained postsynaptic cortical neurons found in the brain, in the thalamus, striatum and red nucleus. The arrow points TdTomato+ neurons. The drawing inset indicates the area corresponding to the image (red square).

D. Absence of TdTomato+ neurons in the deep cerebelar nuclei (of a GAD65-GFP mice injected in the cortex as illustrated in A).

E. In the spinal cord, the labeling is restriced to the descending CS tract (and to its targets) while the ascending tracts are devoided of labeling (CST: corticospinal tract, CC: central canal)

As illustrated in this figure, the targets of the CST are always located in nuclei directly innervated by the cortex, whatever the delay between the injection time and the observation (see also Fig. 4 - Figure supplement 1). The labeling observed in the spinal cord is exclusively derived from anterograde labeling, as transynaptic retrograde transport is not observed in the spinal ascending tracts (nor in the deep cerebelar nuclei). This confirms previous data demonstrating that, when WGA is transgenically encoded (in transgenic mice or by AAVs), the transynaptic labeling is exclusively monosynaptic and anterograde [Libbrecht, S., Van den Haute, C., Malinouskaya, L., Gijsbers, R. & Baekelandt, V. Evaluation of WGA-Cre-dependent topological transgene expression in the rodent brain. *Brain Struct Funct*, doi:10.1007/s00429-016-1241-x (2016); Gradinaru V, Zhang F, Ramakrishnan C, Mattis J, Prakash R, Diester I, Goshen I, Thompson KR, Deisseroth K (2010) Molecular and cellular approaches for diversifying and extending optogenetics. Cell 141:154-165 ; Braz JM, Rico B, Basbaum AI (2002) Transneuronal tracing of diverse CNS circuits by Cre-mediated induction of wheat germ agglutinin in transgenic mice. Proc Natl Acad Sci U S A 99:15148-15153.].

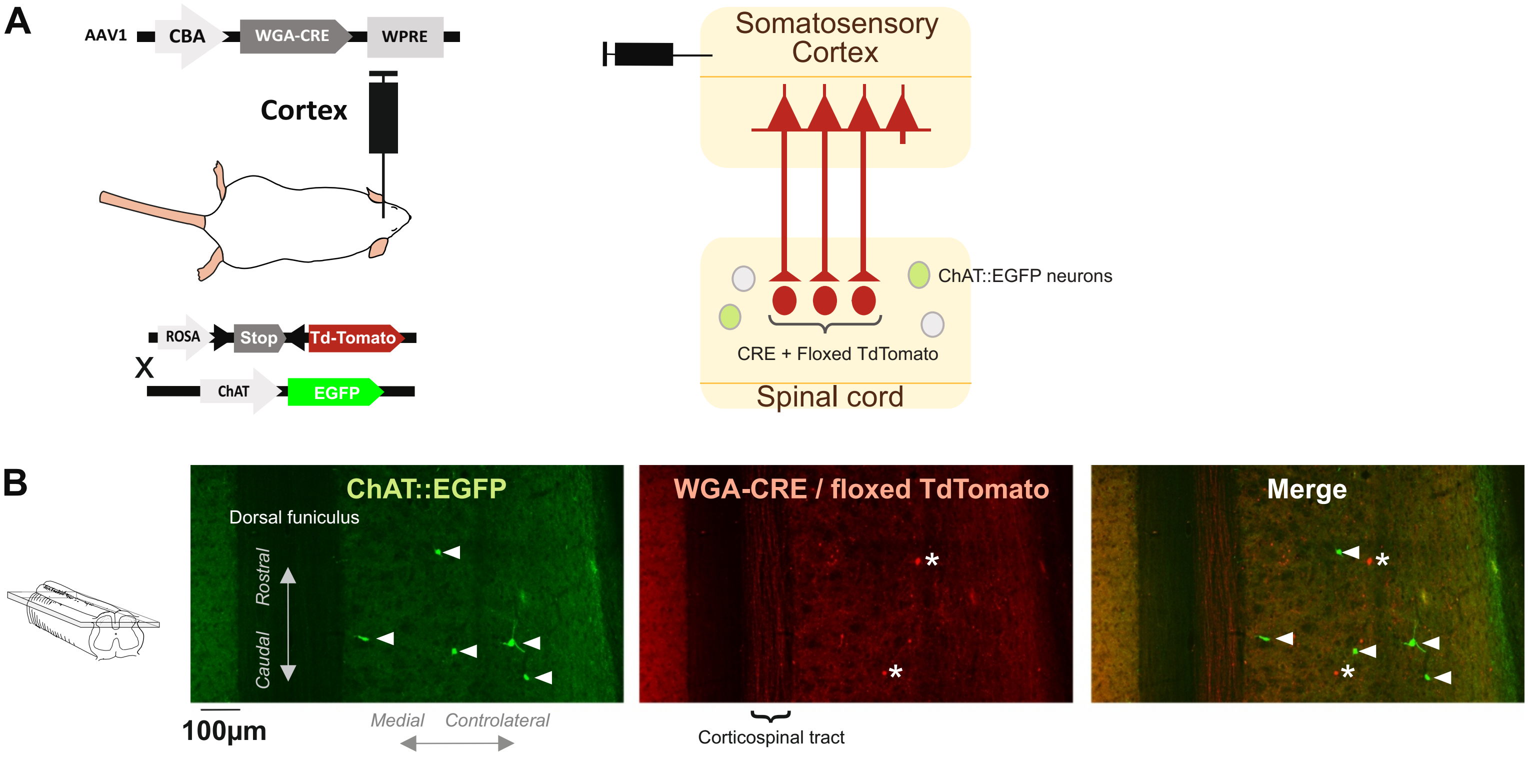


**Figure 1 - Figure Supplement 3. Postsynaptic corticospinal neurons do not colocalize with ChAT.**

A. Schematic experimental design to transynaptically label the targets of the CST in ChAT::GFP transgenic mice. WGA-Cre expressed in the sensorimotor cortex after AAV1 infection can be transfered to the targets of the cortex, for example in the spinal cord ; the mice express a Cre-dependent form of TdTomato, so TdTomato will be expressed in the CST targets after anterograde transynaptic transfer of WGA-Cre. In this experiment the mice are also ChAT::EGFP thus allowing to test whether (some) CST targets are cholinergic.

B. Images from the contralateral dorsal horn of the spinal cord (z-projection of a stack of confocal images, horizontal section located approx. 300µm from the dorsal surface, see inset at the left) showing targets of the CST (TdTomato+, red) and ChAT::GFP+ neurons (green). The arrow heads point at ChAT::EGFP+ neurons located in lamina III-IV, none of which expressed TdTomato. The * points at targets of the CST that do not express ChAT::EGFP.

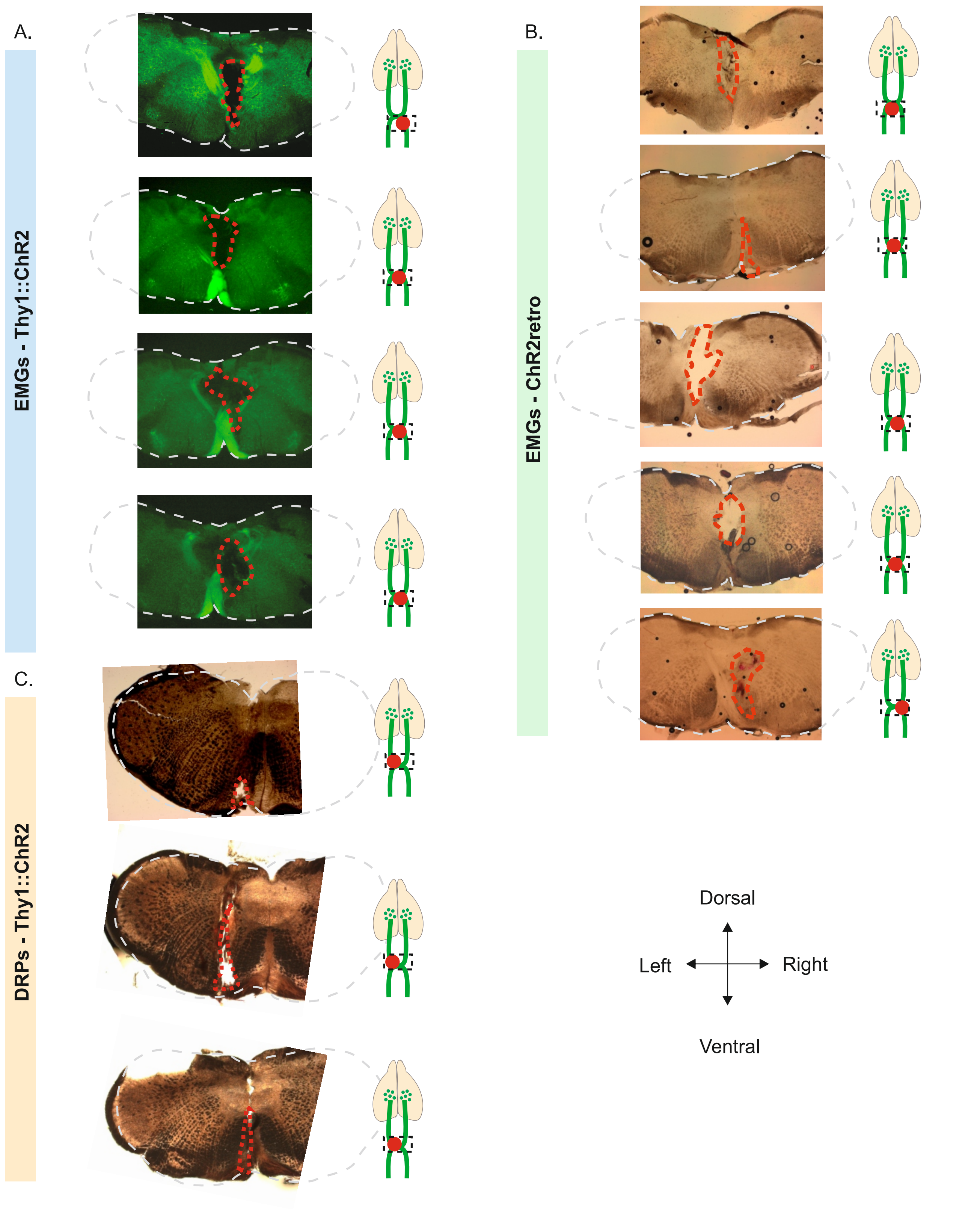


**Figure 2 - Figure Supplement 1. Histology of the pyramidotomies.**
A. Slices of Thy1::ChR2 mice used for EMG recordings at the level of the decussation of the pyramids. Extent of the electrolytic lesion in hatched red.
B. Slices of ChR2 retrogradely labeled mice used for EMG recordings at the level of the decussation of the pyramids. Extent of the electrolytic lesion in hatched red.
C. Slices of Thy1::ChR2 mice used for DRP recordings at the level of the decussation of the pyramids. Extent of the electrolytic lesion in hatched red.

A

| Thy1::ChR2 | 0.5 | 1 | 1.5 | 2 | 2.5 |
|---|---|---|---|---|---|
| **-0.5** | 12.5740682 | 3.37104211 | 19.3783971 | 16.3535276 | 69.863141 |
| **-1** | 7.53966831 | 6.04138632 | 6.80818832 | 35.9359962 | 4.10332195 |
| **-1.5** | 1.38289899 | 8.15556082 | 5.9476861 | 5.92829435 | 2.21480321 |
| **-2** | ND | 9.87158889 | 1.05941526 | 0.53101415 | 4.35257525 |

Z-score > 3
Z-score < 3

B

| Thy1::ChR2 | Mouse 1 | Mouse 2 | Mouse 3 | Mouse 4 |
|---|---|---|---|---|
| **Before Lesion 1** | 7.610458139 | 229.6160926 | 37.53542818 | 21.79331496 |
| **Before Lesion 2** | 5.672072407 | 12.37332975 | 10.20378725 | 41.78392375 |
| **Before Lesion 3** | 5.677845219 | 13.16420671 | 30.07757256 | 43.0112216 |
| **Electrode in decusation** | 22.50949887 | 18.0548327 | 67.66510035 | 103.2101852 |
| **Electrolytic lesion** | 56.50936769 | 28.66464861 | 20.93490606 | 10.76134456 |

C

| ChR2 retroAAV | Mouse 1 | Mouse 2 | Mouse 3 | Mouse 4 |
|---|---|---|---|---|
| **Before Lesion 1** | 16.26824017 | 15.63906278 | 10.3465225 | 30.70771151 |
| **Before Lesion 2** | 6.796548504 | 18.06664921 | 9.457854941 | 4.642069763 |
| **Before Lesion 3** | 10.30119153 | 27.32758053 | 5.189551589 | 15.59097657 |
| **Electrode in decusation** | 2.647111486 | 3.860625737 | -0.61473621 | 1.03036173 |
| **Electrolytic lesion** | -1.103002655 | 1.050864043 | 1.69665654 | 0.681847169 |

D

| GFP retroAAV | Mouse 1 | Mouse 2 | Mouse 3 | Mouse 4 |
|---|---|---|---|---|
| **Spinal photostimulation** | 0.957769 | -0.27026 | 0.130067 | -0.32191 |

E

| ChR2 in CS targets | Mouse 1 | Mouse 2 | Mouse 3 | Mouse 4 |
|---|---|---|---|---|
| **Spinal photostimulation** | -0.219111364 | -0.038196493 | 0.280517075 | 0.001346028 |

Z-score > 1.96
Z-score < 1.96

**Figure 2 - Figure Supplement 2. Z-score of individual EMG responses assessing the significance of the corresponding S/N ratio.**
A. Matrix of the median Z-Score (from 5 animals) for EMG responses recorded at each stimulated point of the cortex. Data presented in Fig. 1C. Coordinates are expressed in mm from Bregma.
B. Z-score of the EMG response for each experimental condition in Thy1::ChR2. Data presented in Fig. 2D.
C. Z-score of the EMG response for each experimental condition in mice injected with the ChETA expressing retroAAV. Data presented in Fig. 3E.
D. Z-score of the EMG responses for each mouse expressing GFP in CS neurons. Data presented in Fig. 3 - Fig. suppl. 1.
E. Z-score of the EMG responses for each mouse expressing ChR2 in the lumbar targets of CS neurons. Data presented in Fig. 4G.

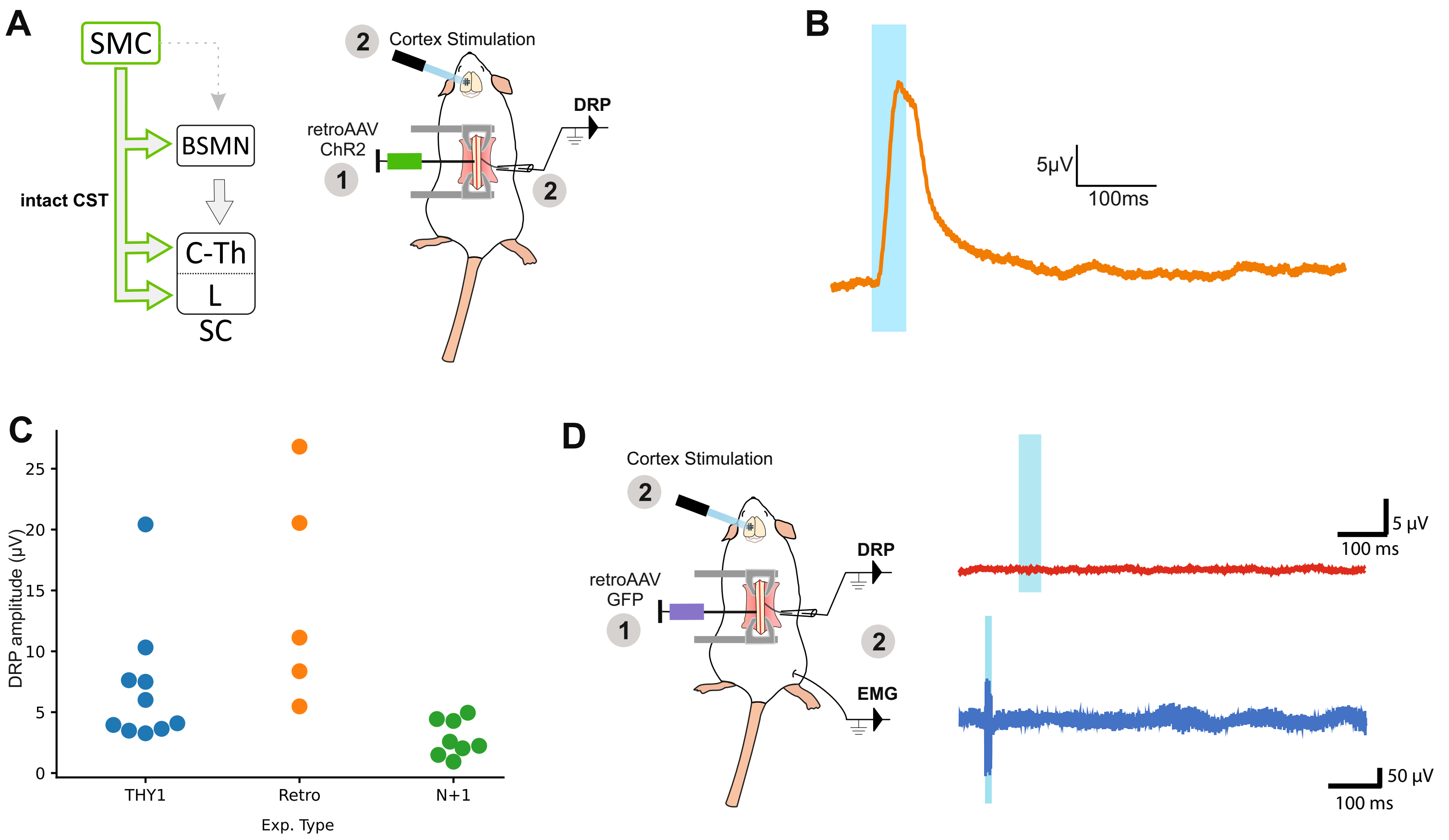


**Figure 3 - Figure Supplement 1. DRPs are induced by photostimulation of ChR2-expressing CS neurons (but not by GFP-expressing CS neurons)**

A. *Left*, diagram illustrating corticospinal neurons and their collaterals expressing ChR2 after retrograde infection in the lumbar cord with a retroAAV. *Right*, experimental design: the photostimulation and DRP recording session took place at least 3 weeks after the infection.

B. Example of DRP recording obtained after cortical photostimulation.

C. Analysis of DRP amplitude in the different animal models : THY1 (recordings presented in Fig. 1 and Fig. 2), Retro (recordings presented in this Figure, panel A-B), N+1 (ChETA expression in the targets of the corticospinal tract, photostimulation of the spinal cord, Fig. 4).

D. *Left*, experimental design of control experiment consisting of the retrograde infection of CS neurons with a control, GFP-encoding, retrograde AAV. *Right*, cortical photostimulation induced no DRP or EMG signal.

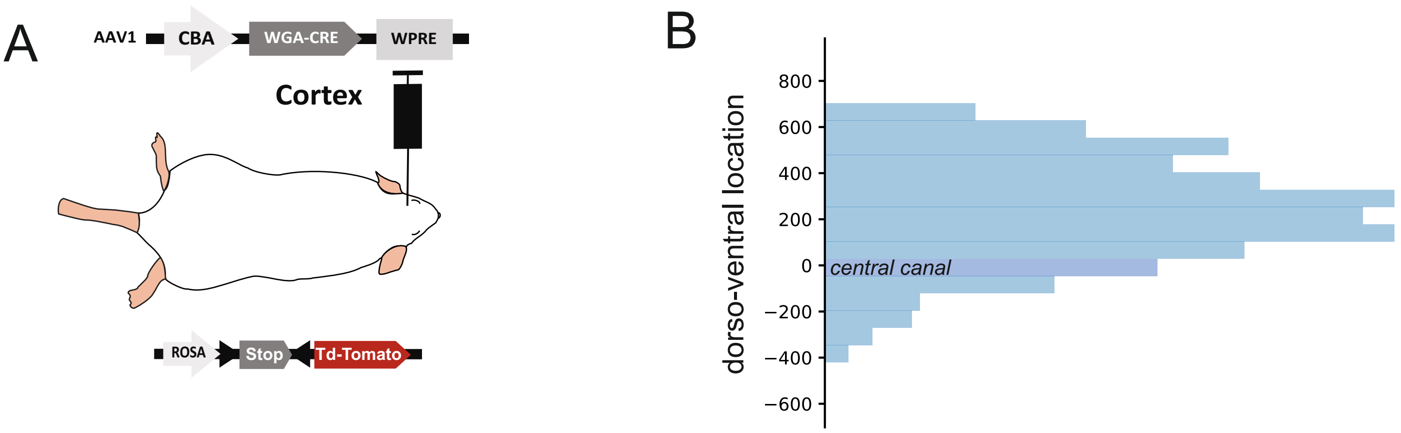


C

| animal ID | Days post injection | nb of spinal TdTomato neurons | Cortical TdTomato area |
|---|---|---|---|
| 8844 | 35 | 53 | 1,375 |
| 8739 | 35 | 55 | 0,725 |
| 8740 | 35 | 229 | 0,544 |
| 4340 | 36 | 6 | 0,046 |
| 8551 | 37 | 14 | 0,294 |
| 7789 | 40 | 11 | 0,159 |
| 4911 | 42 | 4 | 0,728 |
| 5536 | 56 | 10 | 1,207 |
| 6379 | 203 | 188 | 0,991 |

Pearson r 0.5078 0.1922

**Figure 4 - Figure Supplement 1. Strategy to label the spinal targets of corticospinal neurons**

A. Experimental design: the targets of the CST are labeled through a transynaptic approach consisting of AAV1-CBA-WGA-CRE injection in the hindlimb sensorimotor cortex of TdTomato-flex mice.

B. Dorso-ventral distribution of TdTomato neurons in the spinal cord after transynaptic labeling. N=570 neurons from 9 animals. The 0 coordinate corresponds to the center of the central canal. 85% of neurons are located dorsally to the central canal.

C. Number of TdTomato neurons in the spinal cord in the different animals considered for building Fig. 4A, including indication of survival delay after cortical injection, and injection area (in mm²) in the cortex after histological analysis. These parameters do not correlate with the number of Tdtomato neurons as demonstrated by the low Pearson coefficient.

A

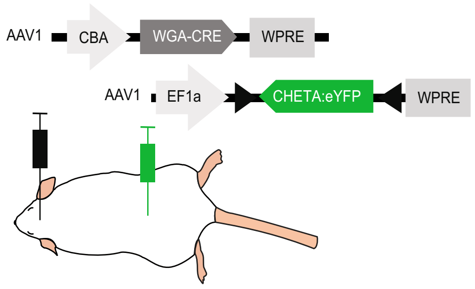


B

| animal ID | nb of ChETA neurons | DRP size (µV) |
|---|---|---|
| 712 | 7 | 4,281 |
| 1181 | 1 | 1,491 |
| 1185 | 18 | 4,435 |
| 2653 | 9 | 2,581 |
| 3105 | 200 | 4,956 |
| 4074 | 6 | 0,000 |
| 3719 | 30 | 0,937 |
| 3723 | 17 | 2,238 |
| 3725 | 14 | 2,035 |

Pearson r 0.17552

C D

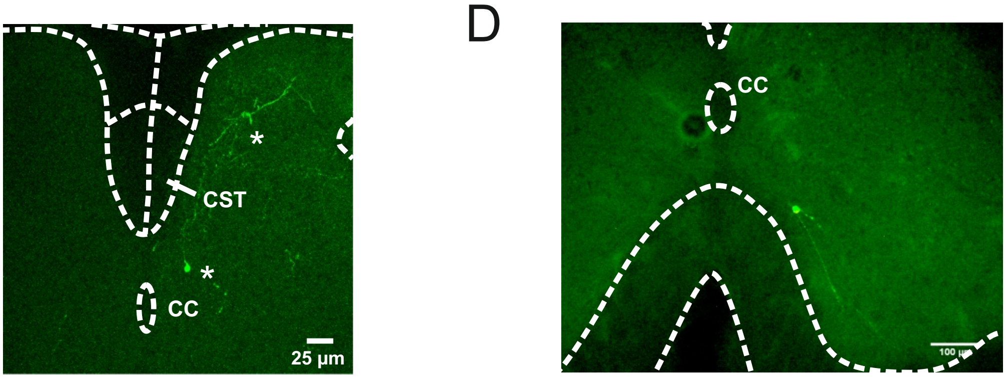


**Figure 4 - Figure Supplement 2. Strategy to stimulate exclusively the spinal targets of corticospinal neurons**

A. Experimental design: the targets of the CST are labeled through a transynaptic approach consisting of AAV1-CBA-WGA-CRE injection in the hindlimb sensorimotor cortex of TdTomato-flex mice.

B. Number of ChETA-expressing targets of the CST, and size of DRP signal recorded in the corresponding mice (no correlation). These numbers be underestimated as the histological analysis was performed after a long recording session, with only post-fixation (no intracardiac perfusion of fixative).

C. After the transectional approach presented in A, the spinal post-synaptic targets of corticospinal neurons express ChETA-eYFP (two *). Importantly the dorsal funiculus, and in particular the corticospinal tract, is not labeled by this approach. CST: corticospinal tract, CC: central canal.

D. The spinal CS post-synaptic neurons (that received the transynaptic WGA-Cre) infected by the intraspinal AAV injection (encoding ChETA-flex) can be located in the ventral horn, as in this example of a neuron located at 180µV ventral to the central canal. Neurons up to -230µV from the central canal were infected, i.e. a depth where 97% of the spinal targets of the CS neurons are found (Figure 4 - Figure supplement 1, panel B)

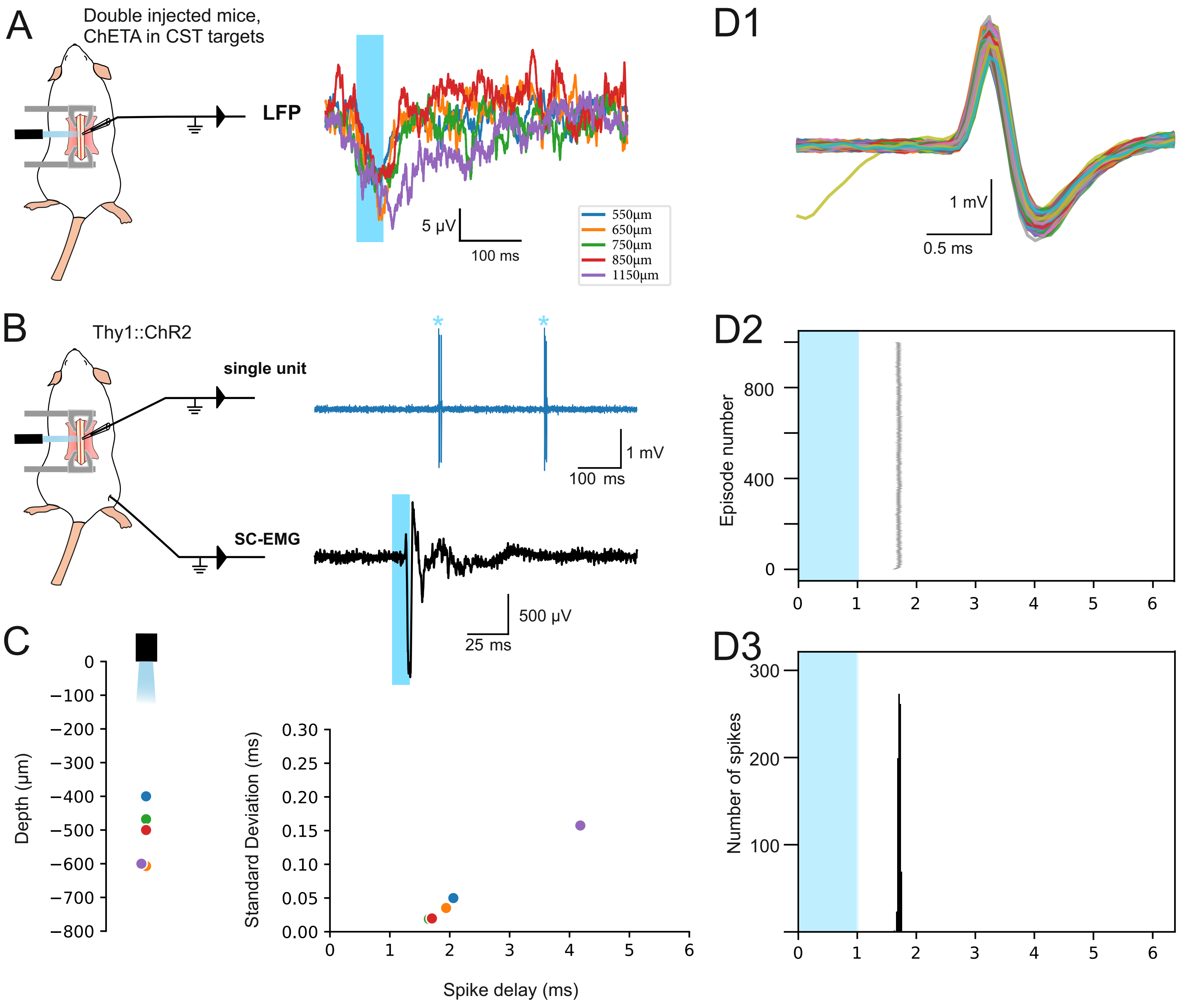


**Figure 4 - Figure Supplement 3. Efficacy of the surface spinal photostimulation**
Control experiments in mice to establish the depth of light penetration after spinal surface illumination

**A.** Experimental design (left) for LFP recordings (right) : the lumbar targets of the CST express ChETA after intersectional viral strategy, and LFP recording are obtained at different depth after spinal surface photostimulation

**B.** Experimental design (left) for spinal single unit (right top) and EMG (right bottom) recordings in Thy1::ChR2 mice, after spinal surface photostimulation (isoflurane anesthesia)
C. (left) Depth of neurons recorded by juxtacellular single-unit recordings, directly responding to the surface illumination (1ms or 0.5ms pulse). (right) Criteria to establish that these neurons were directly activated : the delay of the response was lower than 5 ms, the standard deviation of this delay was lower than 0.20 ms, and (not shown) there was no failure (or less than 0.5% of failure) when stimulated at 4 Hz.
**D.** Example of a spinal neuron located at 500 µm from the cord surface. **D1.** Superimposition of electrophysiological traces of the photostimulated action potential. **D2**. Raster plot of the 1000 episodes illustrating the stability of the response and of its delay. **D3.** Peristimulus histogram presenting the distribution of the spike delay of this neuron in the 1000 episodes presented in D2.

Note : In contrast to the spinal photostimulation presented here, cortical photostimulation of THY1 animals under isoflurane anesthesia did not lead to an EMG signal; this was only obtained under ketamine/xylazine anesthesia.
